# CAMPER: curated annotations for profiling microbial polyphenol metabolic potential

**DOI:** 10.1101/2023.09.24.559193

**Authors:** Bridget B. McGivern, Reed Woyda, Rory M. Flynn, Kelly C. Wrighton

## Abstract

**Summary:** Polyphenols are diverse and abundant carbon sources across ecosystems-having important roles in host-associated and terrestrial systems alike. However, the microbial genes encoding polyphenol metabolic enzymes are poorly represented in commonly used annotation databases, limiting widespread surveying of this metabolism. Here we present CAMPER, a tool that combines custom annotation searches with database-derived searches to both annotate and summarize polyphenol metabolism genes for a wide audience. With CAMPER, users will identify potential polyphenol-active genes and genomes to more broadly understand microbial carbon cycling in their datasets.

**Availability and Implementation:** CAMPER is implemented in Python and is published under the GNU General Public License Version 3. It is available as both a standalone tool and as a database in DRAM v.1.5+. The source code and full documentation is available on GitHub at https://github.com/WrightonLabCSU/CAMPER.

## Introduction

Polyphenols are abundant and diverse plant-derived compounds, spanning 10,000 known chemical structures (Quideau *et al*. 2011). Thus, any environment impacted by plants will be impacted by polyphenols (e.g., soil, human and animal guts) (Hättenschwiler and Vitousek 2000; Cheynier 2005). Despite their abundance and prevalence, polyphenols are often overlooked carbon sources in microbiome studies because the genes encoding polyphenol metabolic enzymes are either missing or poorly organized in existing annotation frameworks. This metabolic blind spot in annotation databases limits the potential for discovery and thorough understanding of microbial carbon cycling.

Annotation, the process of assigning functions to gene sequences, is a crucial step in processing and assigning biological relevance to genomic datasets (Shaffer *et al*. 2020). Most annotation tools use homology-based gene annotations, comparing target sequences to databases of sequences of known-functions. Thus, annotation depends on databases (i) containing broad representation of protein sequences and (ii) containing sufficient amino acid sequence diversity within proteins. However, annotations are only useful if the user can easily infer metabolic context for the annotations.

To overcome these challenges, we present Curated Annotations for Microbial Polyphenol Enzymes and Reactions (CAMPER), a tool that annotates and summarizes microbial polyphenol metabolic potential at scale. CAMPER provides hundreds of expertly-curated annotation searches and summarization at (i) enzyme level recruiting biochemical information where available, (ii) pathway level with data summarized into 102 transformations, and (iii) substrate level with the annotations assigned to the 5 major classes of polyphenols.

## 2) CAMPER Development and Usage

The CAMPER workflow is outlined in **Figure 1A-B**.

**Figure 1.**
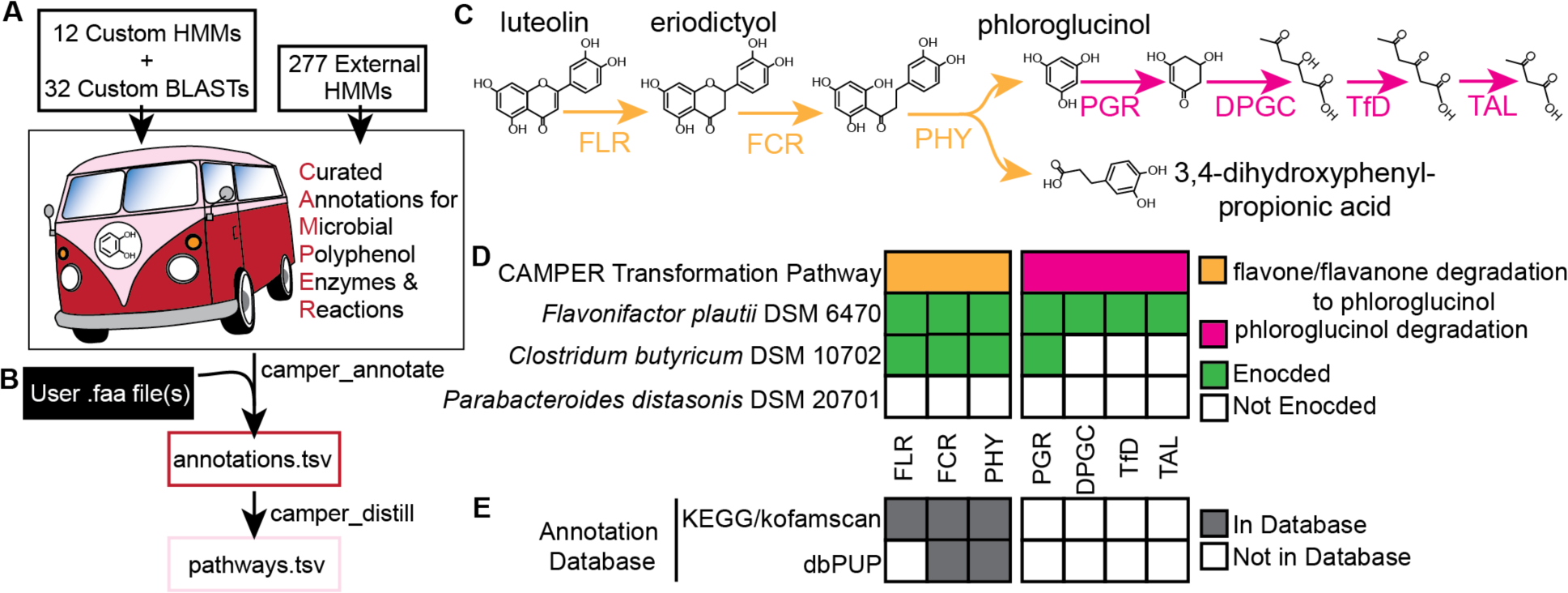
CAMPER workflow and application. (**A**) CAMPER annotation library is comprised of 12 custom hidden Markov models (HMMs), 32 custom Basic Local Alignment Search Tool (BLAST) searches, and 277 HMMs from other databases. (**B**) CAMPER takes as input one or multiple amino acid FASTA files, and using *camper_annotate* generates an annotations.tsv file and through *camper_distill* generates a summarized tsv file. (**C**) Examples of two CAMPER transformation pathways, Flavone/Flavanone degradation to phloroglucinol (orange arrows) and phloroglucinol degradation (pink arrows). (**D**) CAMPER annotated presence for seven genes in three isolate genomes, where green marks the presence of an A or B rank annotation. (**E**) The presence of the 7 example enzymes in other common or polyphenol-focused annotation databases, where grey indicates presence in the database.

### 2.1 CAMPER Annotation Library Development

CAMPER provides a library of custom annotations, including 12 HMMs and 32 BLAST searches (**Fig. 1A**). To build the custom annotation searches, we identified 44 biochemically-characterized polyphenol-active enzymes that were either absent or poorly described in commonly-used annotation datasets (**Supplementary Table 1**). Importantly, these enzymes come from microorganisms from diverse ecosystems, making the suite of pathways relevant to users across microbiomes.

If more than three sequences existed for an enzyme, and were representative of more than one phylum, we constructed an HMM for that sequence similar to in (Khot *et al*. 2022). Briefly, experimentally validated “seed” sequences and decoy sequences (ie. sequences from closely related but biochemically distinct functions) were BLAST searched against the UniRef90 database (August 2021)(Suzek *et al*. 2007), and the top 300 hits for each sequence were pulled. Then the seed, decoy, and UniRef90 hit sequences were aligned with MAFFT (v7.055b) (Katoh and Standley 2013) using the “-auto” flag, and the alignment was trimmed using trimal (v1.4.rev22) with the “-gappyout” flag (Capella-Gutierrez, Silla-Martinez and Gabaldon 2009). The trimmed alignment was used to construct a tree using IQ-Tree (v1.6.8) (Nguyen *et al*. 2015) with the following flags: -alrt 1000 -bb 1000 -m MFP -nt AUTO -ntmax 10. The tree was visualized in iTOL (Letunic and Bork 2021), and rooted on the clade containing all decoy sequences. Then, we identified the monophyletic clade containing all seed sequences, and pulled these sequences to use for HMM construction. Importantly, we removed the seed sequences from this set. The remaining non-seed sequences were used to construct an HMM using graftM create (v.0.13.0) (Boyd, Woodcroft and Tyson 2018). The HMM was then searched against the full set of sequences (seed, decoy, UniRef90 hits) using graftM graft.

We then assigned two bitscore thresholds for each HMM (**Supplementary Table 2**): “A” scores were greater than or equal to the lowest bitscore obtained from aligning the HMM against the experimentally validated seed sequences, representing high-confidence annotations. “B” scores corresponded to the lowest bitscore obtained from aligning the HMM against sequences that form monophyletic clades with the seed sequences. Notably, the “B” scores were higher than the bitscore obtained from aligning the HMM against decoys, thus they represent exploratory annotations.

When there was not sufficient numbers of, or diversity in, sequences available for an enzyme, we used a BLAST approach using the validated seed sequence. We used an “A” bitscore cutoff of 200 and a “B” cutoff of 120 for the BLAST searches. We combined these custom BLAST and HMM annotations with 272 HMMs from kofamscan (Aramaki *et al*. 2020) and 5 HMMs from dbCAN2 (Zhang *et al*. 2018), using the recommended score cut-offs provided by those annotation tools.

### 2.2 Running CAMPER

CAMPER can be ran as a standalone tool (CAMPER github), or it can be used as a database within DRAM (Shaffer *et al*. 2020) (from v1.4.4 on) using the “--use-camper” flag. In the standalone tool, there are two steps: *camper_annotate* and *camper_distill*. The first step, *camper_annotate*, takes as input amino acid sequences in FASTA format (**Fig. 1B**). This can be a single input file (ex. one genome, one assembly, concatenated genomes), or a wildcard pointing to many files (ex. multiple genomes, multiple assemblies). This step searches the CAMPER annotation library against the input sequences using hmmscan (HMMER v3.3) (Eddy 2011) or blastp (blast v2.8.1) (Camacho *et al*. 2009), parses the search results, and outputs *annotations.tsv*, a tab-delimited file where each row corresponds to an input gene sequence, and the columns provide information for A and B scoring hits (**Fig. 1B, Supplementary Table 3**). Then, the *camper_distill* step takes this *annotations.tsv* file as input, and outputs a user-named tab-delimited file with the polyphenol context for each gene and the counts of those genes in each input file (columns) (**Fig. 1B, Supplementary Table 4**).

### 2.3 CAMPER Outputs

The distilled CAMPER output file provides curated context for genes encoding polyphenol-active enzymes (**Fig. 1C**). First, annotations are organized at the gene level and subsequently into transformation pathways. For each transformation pathway, the type of polyphenol used as substrate is noted in a hierarchical structure according to Phenol-Explorer ontology (Rothwell *et al*. 2013). Additionally, the oxygen requirement for each pathway is given: “anoxic” transformations were uncovered in anoxic ecosystems (ex. human gut) or demonstrated to act under anoxic conditions, while “oxic” transformations were demonstrated to act under oxic conditions or directly require oxygen (ex. dioxygenases). Some transformations are noted as “both” because they are inferred to operate under both conditions (ex. tannase). Finally, the specific reaction catalyzed in the step is provided. Then, the gene counts of each annotation is given for each input file, where each input file is a column (**Supplementary Table 4**). CAMPER contains 321 annotations, organized into 102 transformation pathways (**Supplementary Figure 1**). We have also provided a key for quickly identifying pathways for specific polyphenols of interest (**Supplementary Table 5**).

## 3) CAMPER rapidly and accurately identifies genes for flavanone degradation in microbial genomes

To illustrate the ability of CAMPER to rapidly profile and summarize microbial polyphenol metabolic potential, we annotated genomes from known and hypothesized polyphenol degrading microorganisms. Here, we focus on two anoxic CAMPER transformation pathways spanning 7 enzymes (**Fig. 1D**): (i) flavone/flavonone degradation to phloroglucinol, which encompasses the enzymes flavone/flavonol reductase (FLR), flavanone/flavanonol-cleaving reductase (FCR), and phloretin hydrolase (PHY), and (ii) phloroglucinol degradation, covering enzymes phloroglucinol reductase (PGR), dihydrophloroglucinol cyclohydrolase (DPGC), triacetate-forming dehydrogenase (TfD), and triacetate acetoacetate-lyase (TAL).

The first genome we annotated was from *Flavonifractor plautii* DSM 6470, a model organism for polyphenol degradation. *F. plautii* is known to degrade flavonoids and phloroglucinol, and the enzymes it uses have been characterized (Braune and Blaut 2016). In fact, the *F. plautii* genes were used as seed sequences in building the HMMs for all 7 enzymes. As expected, CAMPER gives “A” rank annotations for all 7 genes (**Fig. 1E, Supplementary Table 3)**. Thus, CAMPER can reliably annotate known genes.

Next, we used CAMPER to annotate genes in non-model microorganisms. To do this, we selected a genome for *Clostridium butyricum* DSM 10702, an isolate that was experimentally shown to degrade the flavanone eriodictyol to phloroglucinol and 3,4-dihydroxypropionic acid (Miyake, Yamamoto and Osawa 1997) (**Fig. 1D**). Importantly, the underlying flavanone degradation genes and enzymes in *C. butryicum* have not been identified. Using CAMPER, we identified B and A rank annotations for FCR and PHY, respectively, providing candidate genes to support the phenotype (**Fig. 1D-E, Supplementary Tables 3-4**). In the original study, the degradation product phloroglucinol was not detected in the medium of *C. butryicum* incubated with eriodictyol, suggesting it was rapidly consumed (Miyake, Yamamoto and Osawa 1997). In support of this, *C. butryicum* encodes a gene annotated with a B-rank as PGR (**Fig. 1E, Supplementary Table 3**). We did not obtain A or B rank gene annotations for the rest of the phloroglucinol pathway, indicating *C. butyricum* either encodes an alternative cryptic transformation pathway, or it only catalyzes the initial reduction step. Beyond phloroglucinol, *C. butyricum* encodes a gene annotated with an A rank as FLR, suggesting it may also be able to carry out an upstream reaction converting the flavone luteolin to eriodictyol (**Fig. 1D-E, Supplementary Tables 3-4**). Collectively, this example highlights the role CAMPER can play in both testing and generating hypotheses.

Finally, we annotated a third genome for *Parabacteroides distasonis* DSM 20701, noted in the study by Miyake et al. for not being able to degrade eriodictyol. Mirroring this laboratory phenotype, CAMPER did not annotate any genes as FCR or PHY (**Fig. 1E, Supplementary Table 3**). Therefore, CAMPER’s score thresholds appear to control for false-positive annotations.

These three genomes are a case study in the ability of CAMPER to identify and easily summarize genes for polyphenol metabolism in both well-studied and under-explored genomes, as well as control for false positive annotations. To obtain comparable data using existing annotation databases, we tried to annotate the same three genomes with kofamscan and dbPUP (Zheng *et al*. 2022), another polyphenol-focused annotation database. Of the 7 enzymes, kofamscan and dbPUP included 3 and 2, respectively, in their database (**Fig. 1F**). In fact, both were missing annotations for the entire phloroglucinol degradation pathway, demonstrating the value that CAMPER brings for curated pathways.

Practically, it took 57 seconds to run *camper_annotate* when giving the three genomes as input at once, and 2 seconds to run *camper_distill*. To annotate with dbPUP, the genomes had to be downloaded and then uploaded to the dbPUP web interface, complicated by the fact that there was not a recommended score cut-off for the searches and all annotations are reported at the same level. From both databases, the results needed to be manually curated to gain pathway context, creating a knowledge barrier to widespread use. CAMPER was designed to overcome these hurdles, providing rapid and thorough summarization of the annotations.

## 4) Conclusion

Here we present CAMPER as a fast and user-friendly tool for identifying polyphenol metabolism in microbial genomic datasets. CAMPER can be run as a standalone tool or be used as a database in the annotation software tool Distilled Refined Annotation of Metabolism (DRAM). We showed that CAMPER can identify polyphenol metabolic pathways in well-studied and suspected polyphenol metabolizing genomes, giving results that mirror experimental evidence. CAMPER makes microbial polyphenol metabolism accessible to a wide audience, facilitating discovery and a better understanding of microbial carbon cycling.

