## Supplementary figures and images for "CAMPER: curated annotations for profiling microbial polyphenol metabolic potential"

### Supplementary Figure 1

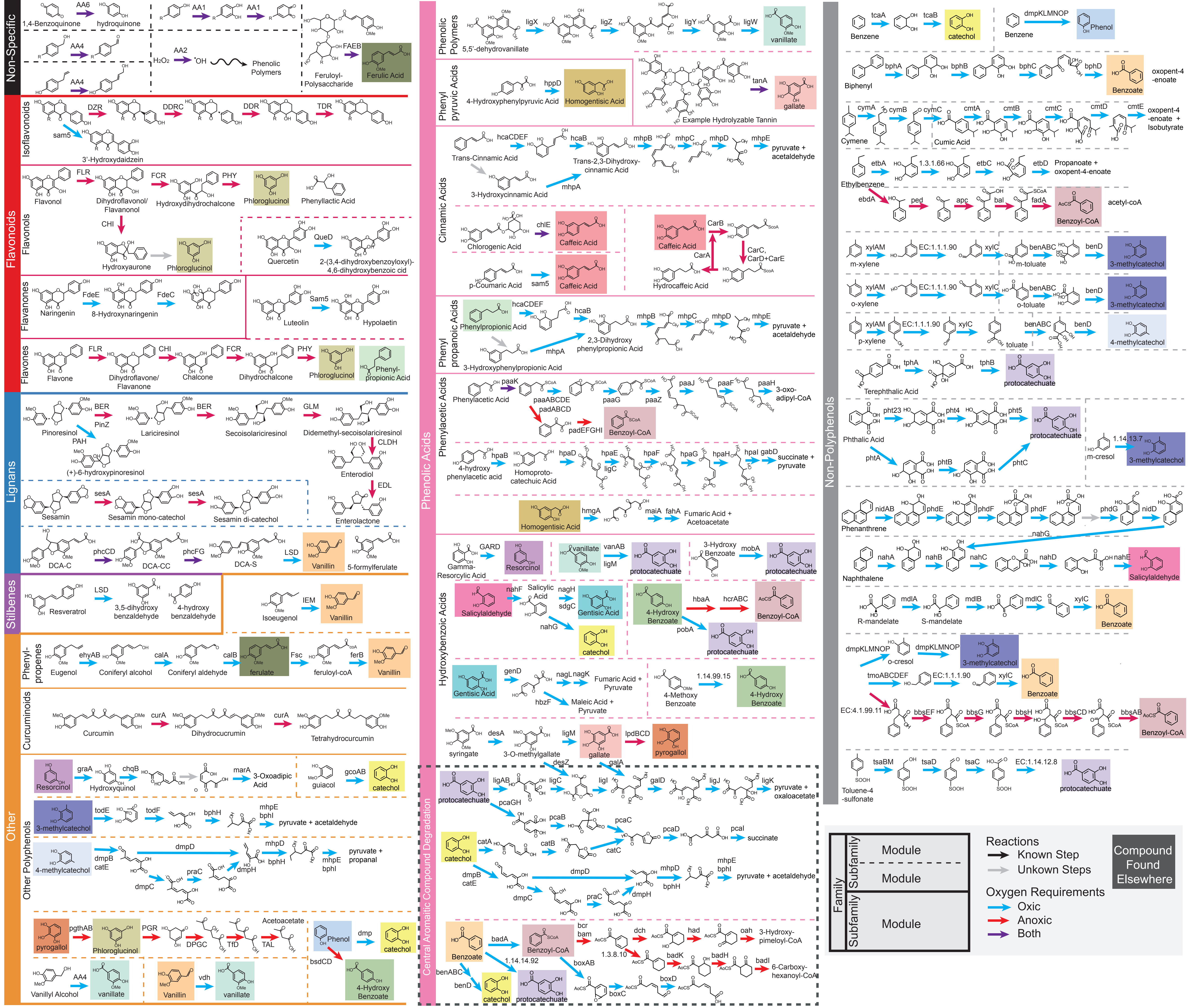
